# Increased bleeding and thrombosis in myeloproliferative neoplasms mediated through altered expression of inherited platelet disorder genes

**DOI:** 10.1101/2023.05.23.541977

**Authors:** Alan Mitchell, Mattia Frontini, Serajul Islam, Suthesh Sivapalaratnam, Anandi Krishnan

## Abstract

An altered thrombo-hemorrhagic profile has long been observed in patients with myeloproliferative neoplasms (MPNs). We hypothesized that this observed clinical phenotype may result from altered expression of genes known to harbor genetic variants in bleeding, thrombotic, or platelet disorders. Here, we identify 32 genes from a clinically validated gene panel that were also significantly differentially expressed in platelets from MPN patients as opposed to healthy donors.

This work begins to unravel previously unclear mechanisms underlying an important clinical reality in MPNs. Knowledge of altered platelet gene expression in MPN thrombosis/bleeding diathesis opens opportunities to advance clinical care by: (1) enabling risk stratification, in particular, for patients undergoing invasive procedures, and (2) facilitating tailoring of treatment strategies for those at highest risk, for example, in the form of antifibrinolytics, desmopressin or platelet transfusions (not current routine practice). Marker genes identified in this work may also enable prioritization of candidates in future MPN mechanistic as well as outcome studies.

## Introduction

Myeloproliferative neoplasms (MPNs) are associated with increased bleeding and thrombotic events^1-6^, that are also a significant cause of morbidity and mortality^7-9^ in these patients. Alongside molecular tests, bone marrow biopsy is a cornerstone of MPN diagnosis^10,11^, with subtype determination informed by characteristic cell morphology and fibrosis assessment. Yet this procedure is not without complication for some patients; and apparent risk factors for hemorrhage post bone marrow aspiration include myeloproliferative disorders.

However, the mechanisms underlying MPN association with thrombo-hemorrhagic events remain unclear^6,12,13^. It has previously been suggested that in MF, hemorrhagic events result from a combination of progressive thrombocytopenia, secondary to bone marrow failure, and abnormalities in platelet function^14,15^.

We therefore hypothesized that platelets in MF would demonstrate altered expression of a specific set of genes that are known to be associated with inherited bleeding, thrombosis, and platelet disorders, as characterized (and continuously under review) by an expert panel^14,16^, and already validated for clinical use^17-19^. Here, we interrogate this specific gene set for differential platelet expression in a comprehensive cohort of platelet transcriptomes^20^ from patients with chronic progressive MPNs.

## Methods

Study approval was provided by the Stanford University Institutional Review Board (#18329). We collected blood from MPN patients enrolled in the Stanford University and Stanford Cancer Institute Hematology Tissue Bank after written informed consent from patients or their legally authorized representative. Eligibility criteria included age ≥18 years and Stanford MPN clinic diagnosis of essential thrombocythemia, polycythemia vera or myelofibrosis (defined using the consensus criteria at the time of this study). We use the term ‘myelofibrosis’ to encompass both primary myelofibrosis and myelofibrosis evolved from essential thrombocythemia or polycythemia vera. For healthy controls, blood was collected from adult donors selected at random from the Stanford Blood Center. All donors were asked for consent for genetic research. Altogether, our platelet transcriptome dataset^20^ comprised 118 human peripheral blood samples as follows: healthy controls (n=21) and World Health Organization-defined MPN patients (24 ET, 33 PV and 40 MF) including seven untreated, and 92 either on cytoreductives/biologics (*e*.*g*. ruxolitinib, hydroxyurea, interferon-alpha), anti-thrombotic agents (e.g. aspirin, warfarin), or a combination of these reflecting the diversity among MPN patients.

### Platelet isolation, library preparation, and RNA sequencing

All blood samples were collected into acid citrate-dextrose (ACD, 3.2%) sterile tubes (Becton, Dickinson and Co.) and platelets were isolated with an established protocol^21-24^ within 4 h of collection. For RNA-sequencing (RNA-seq), 1×10^9^ isolated platelets lysed in Trizol were processed to extract RNA (all integrity numbers >7.0) and library preparation. Twelve pooled samples with individual indices were run on an Illumina HiSeq 4000 (Patterned flow cell with Hiseq4000 SBS v3 chemistry) as 2 × 75bp paired end sequencing with a coverage target of 40M reads/sample.

### Statistical Analysis and Variant-Expression Mapping

Platelet transcriptomic data were library-size-corrected, variance-stabilized, and log2-transformed using the R package DESeq2^25^. The same algorithm was used to determine differential gene expression, while adjusting for patient age, gender and treatment as confounding variables and controlling for multiple comparisons using the Benjamini-Hochberg defined false discovery rate (FDR). Significant variance in expressed transcripts were pre-specified as transcripts with an FDR <0.05 and a log2 fold change ≥ 0.5 in MPN, as compared to healthy controls (the entire differential transcriptome was applied toward downstream Gene Set Enrichment Analysis).

Using Mann-Kendall trend test (multiple comparisons adjusted with the Benjamini-Hochberg method) on normalized gene counts, we assessed progressive and monotonic upward or downward trends in gene expression and identified statistically significant progressive genes across the three MPN subtypes, ET, PV, and MF at a false discovery rate FDR <0.05.

## Results and Discussion

The bleeding, thrombotic, and platelet disorder panel^17,26^ constituted a total of 145 unique genes. Of these, 32 were found to be progressively differentially expressed (FDR <0.05) across MPNs. **Table 1** details these genes with results from both pairwise differential comparisons (each MPN subtype versus healthy donors) as well as progressive expression trend analysis across all three MPN subtypes.

**Table 1:**
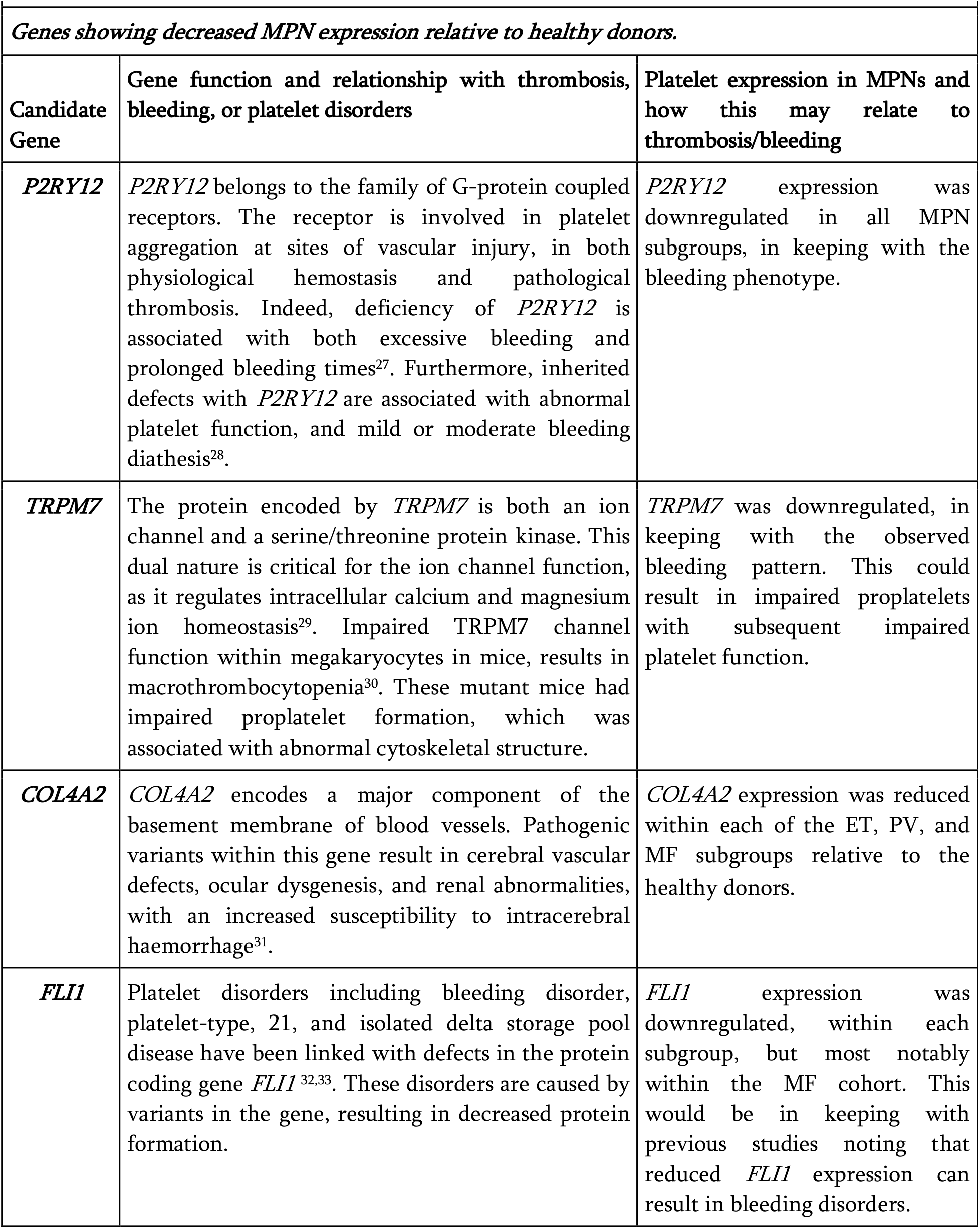
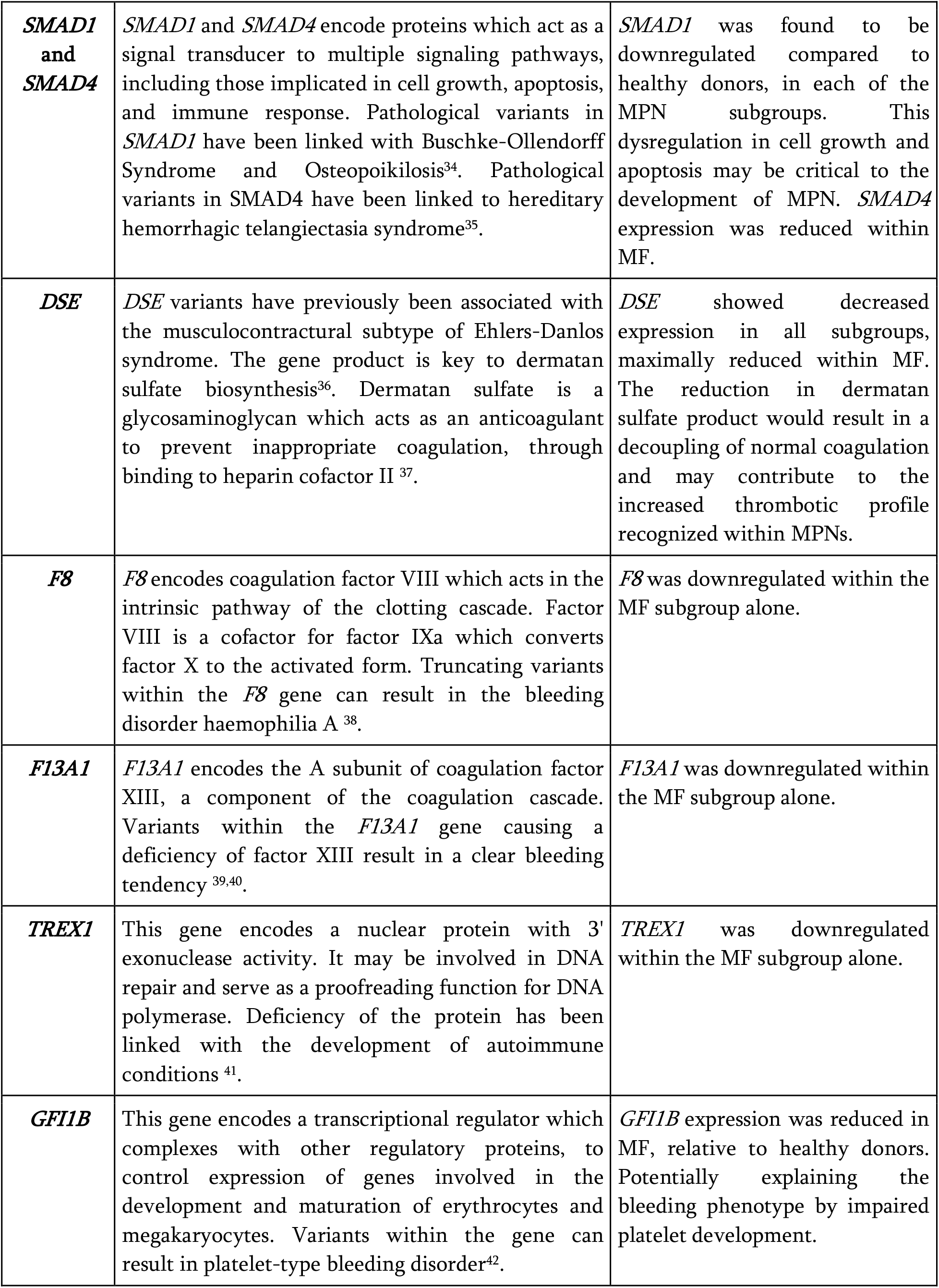
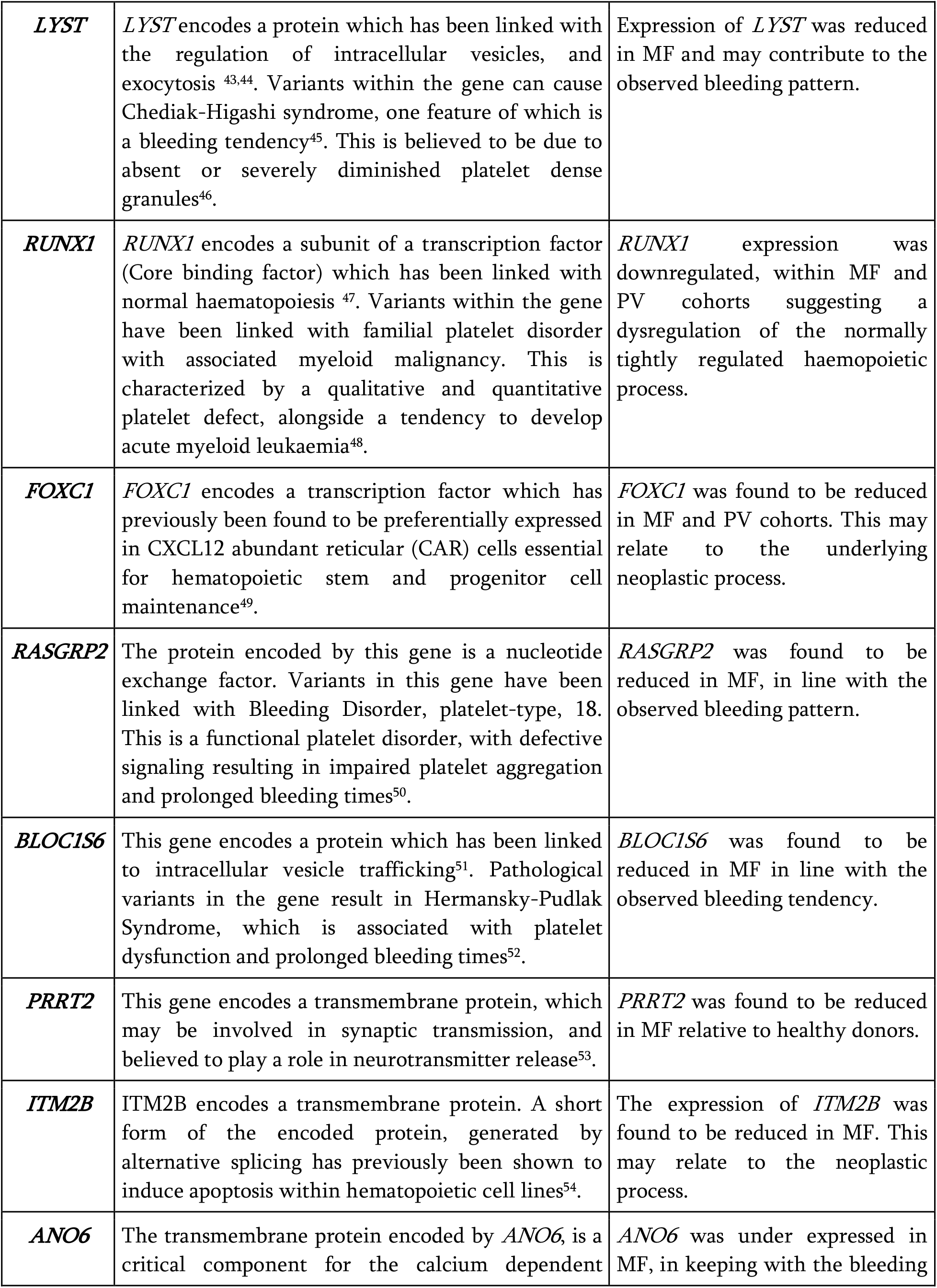
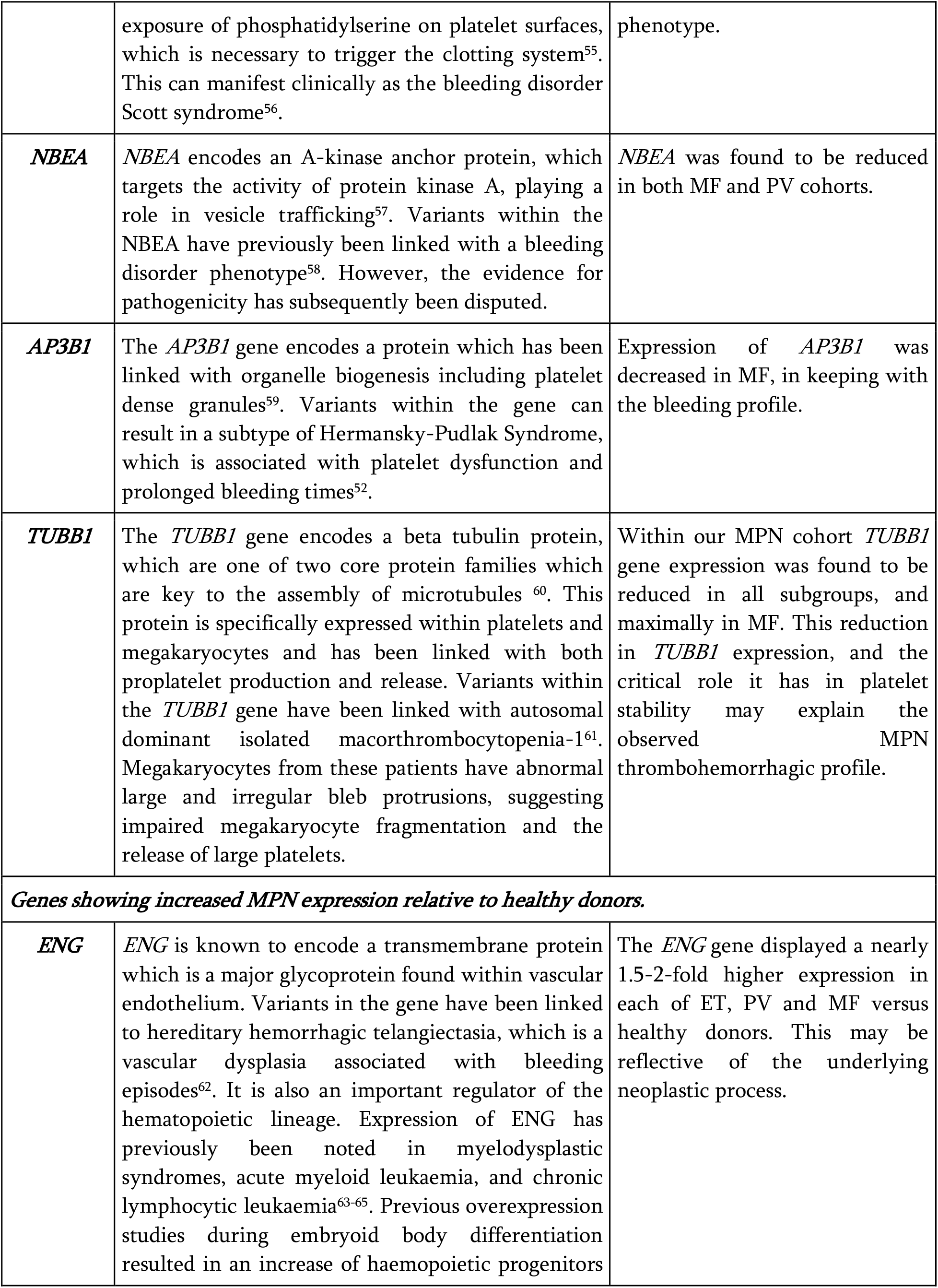
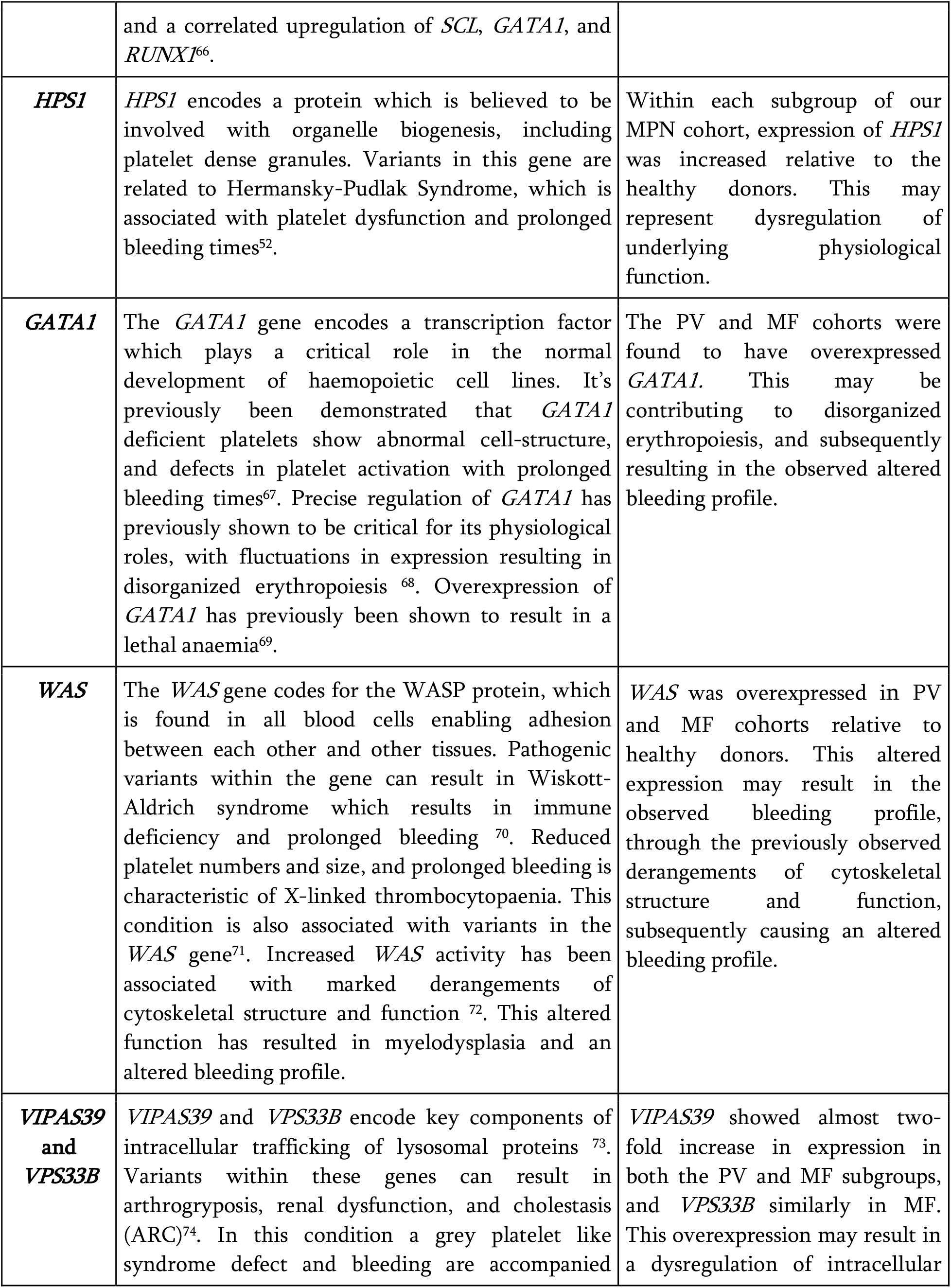
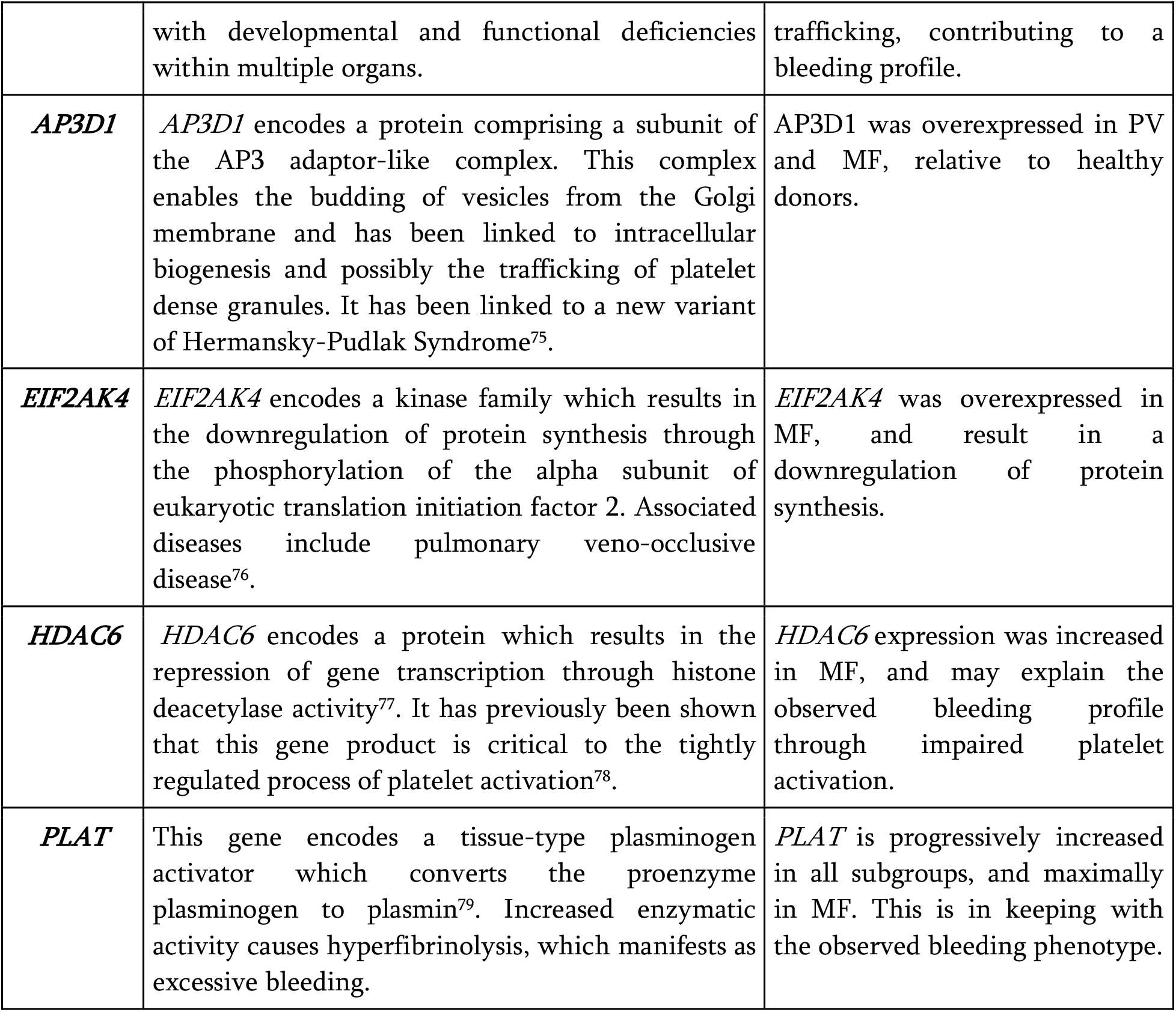
Inherited bleeding and platelet disorder genes whose altered expression may explain the increased thrombo-haemorrhagic risk in myeloproliferative neoplasms. Column 1 identifies the candidate gene of interest, Column 2 provides a summary of gene function, and evidence of relationship with either platelet or bleeding disorders and Column 3 highlights the observed differential either as a pairwise comparison between any given MPN subtype versus healthy donors or as a progressive change across all three subtypes.

The top 12 progressively differentially expressed genes are also visualized in **Figure 1**. 22 genes showed reduced expression in MPNs relative to healthy donors. These included genes previously linked with granule release and development such as *AP3B1, LYST*, and *BLOCS6*, and disorders of platelet function such as *P2RY12* and *AN06*. One noteworthy example is the discordant expression of known anticoagulant genes (*e*.*g. DSE* downregulation) versus those of platelet dense granule function (*e*.*g. HPS1* upregulation) across MPNs, likely a factor in the uncertain risk of both thrombosis and bleeding in MPNs. 10 genes were overexpressed compared to healthy donors. Some of the increased expression levels may reflect the underlying neoplastic process in MPNs, such as that observed for *ENG and RUNX1*. Others, such as *PLAT, GATA1* and *HDAC6* have been linked to impaired platelet development or activation and may contribute to the bleeding profile observed in MPNs.

**Figure 1:**
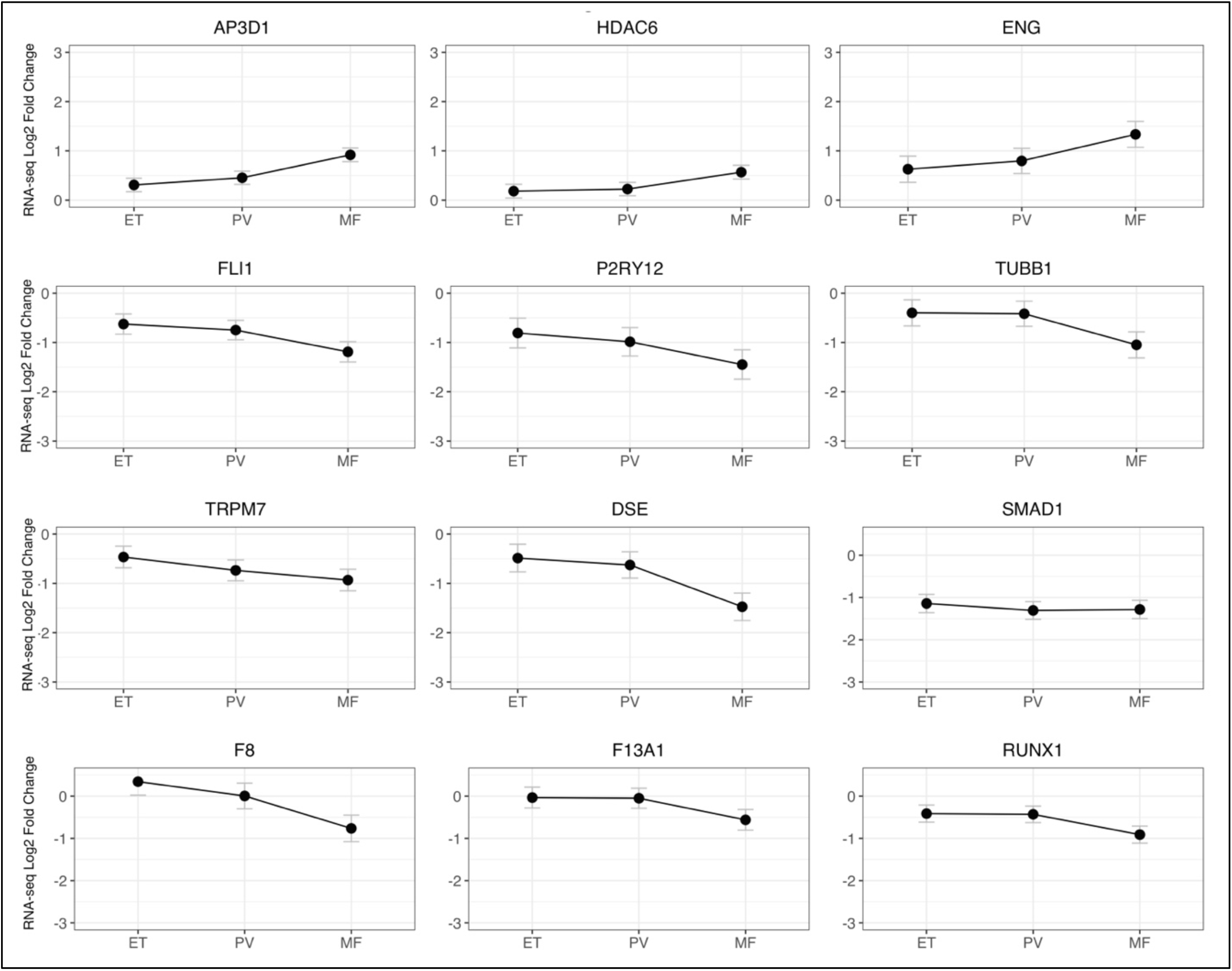
Progressive expression of markers of thrombohemorrhagic risk in MPNs. Top 12 genes (out of 32 detailed in Table 1) demonstrating monotonic progressive gene expression (log2 fold change in expression *y*-axis, FDR < 0.01, *Mann-Kendall* test with Bonferroni correction) across *x-axis* MPN subtypes (ET/PV/MF) versus healthy donors (CTRL).

Together, these genes offer opportunities for evaluation, not only as biomarkers in assessment of risk, prognosis and monitoring for thrombosis/bleeding in MPN patients but also as candidates for mechanistic interrogations in MPN model systems and future outcome studies.

## Author Contributions

A. Krishnan, M. Frontini and S. Sivapalaratnam conceived of the overall study. A.M. and A.K. wrote and edited the manuscript. M.F., A.K. and S.S. critically reviewed and edited the manuscript. All authors approved the final manuscript.

## Acknowledgements

This work was funded by US National Institutes of Health grants 1K08HG010061-01A1 and 3UL1TR001085-04S1 and the MPN Research Foundation Challenge Grant to A.K, the UK National Institute for Health Research to A.M and S.S., and the British Heart Foundation Senior Basic Research Fellowship (FS/18/53/33863) to M.F. Authors thank Dr. Jason Gotlib at Stanford University, the MPN patients at the Stanford Cancer Institute, and the healthy donors at the Stanford Blood Center for their contribution to this research. A.K. extends special thanks to Dr. Sarah Kelliher (ISTH Training Fellow at Stanford from University College Dublin) for her critical reading of this manuscript.

